# Modelling the polygenicity and clinical heterogeneity of human depression in mice to identify biomarkers of antidepressant response

**DOI:** 10.64898/2026.03.31.715499

**Authors:** Claire Altersitz, Sébastien Arthaud, Martine Dubois, Violaine Latapie, Jean-Marie Vaugeois, Malika El Yacoubi, Stéphane Jamain

**Author notes:** <u>Corresponding authors:</u> Dr Stéphane Jamain, Inserm U955, Neuropsychiatrie Translationnelle, Faculté de Santé, Université Paris Est Créteil, 8 rue du Général Sarrail, Créteil, 94000, France., Dr Malika El Yacoubi, Inserm U955, Neuropsychiatrie Translationnelle, Faculté de Santé, Université Paris Est Créteil, 8 rue du Général Sarrail, Créteil, 94000, France. Contributed equally (Jean-Marie Vaugeois, Malika El Yacoubi, Stéphane Jamain).

## Abstract

Major depressive disorders (MDD) are predicted to become the first cause of burden of disease worldwide in 2030, but 30% of patients still do not respond to antidepressants. Current rodent models of MDD mainly result either from one genetic or one environmental risk factor exposure, not recapitulating the multifactorial and polygenic nature of MDD. We recently generated a polygenic mouse model of MDD from selective breeding after mild stress in the Tail Suspension Test (TST), named H-TST. Here, we selected animals exhibiting high immobility during the Forced Swim Test (FST) to generate a new stable polygenic model of MDD, called H-FST. Unlike our previous H-TST model, H-FST mice did not exhibit any anxiety-or anhedonia-like behaviors, nor did they display any sleep disturbances. Moreover, H-TST and H-FST mice showed opposite response after administration of various antidepressant treatments. The gene expression level in the prefrontal cortex of H-TST and H-FST mice revealed little overlap in genes and biological pathways associated with depressive-like behaviors and opposite dysregulation of excitatory/inhibitory synaptic imbalance. Finally, these two models allowed in humans the identification biomarkers of treatment response specific of clinical subgroup of patients.

## INTRODUCTION

Major depressive disorders (MDD) affect 280 million people worldwide, twice as many women as men, and are among the most prevalent mental health conditions.^1^ It is also a major contributor to the overall global burden of disease.^1^ Treatments are only partially effective in affected individuals and there are currently no clinical signs or biomarkers that can predict improvement in patients’ symptoms. This illustrates our lack of knowledge on the etiopathology of the disease.^2–4^ Genome-wide association studies in large cohorts have identified hundreds of genes associated with an increased risk of developing MDD.^5^ Although animal models are of paramount importance in understanding the pathophysiology and identifying biological markers, this polygenic vulnerability is difficult to model in rodents. To date, available mouse models developed to decipher MDD pathogenesis focused either on the contribution of a single gene or a single environmental risk factor and do not recapitulate the multifactorial etiology of MDD.^6^ To address this issue, we developed in the past two models of depressive-like behaviors, named H/Rouen and H-TST, by selecting mice for their passive coping mechanism, so called helplessness, when facing a stressful situation in the tail suspension test (TST), a reference test used to assess the efficacy of antidepressants.^7–9^ H/Rouen and H-TST mice showed a near absence of mobility in the TST unlike their non-helpless counterparts, NH/Rouen and NH-TST, which showed a persistence in the escape behavior throughout the test. Note that these behaviors were stable through generations and associated with other depressive-like behaviors, including anhedonia-like, anxiety-like behaviors, and sleep impairments.^7–11^ Exome sequencing identified several hundred genes commonly associated with helplessness in these models, a majority of them have also been associated with MDD in humans.^9^ Beside the TST, the forced swim test (FST) is the second most commonly used test to assess antidepressant efficacy.^12^ It is unclear whether these two tests produce stress through shared mechanisms, but they have been clearly shown to provide variation in antidepressant response.^13–15^ Here, we developed a new mouse model of MDD based on bidirectional selection of extreme phenotypes in the FST and starting from the same complex genetic background used to generate H-TST and NH-TST mice. This selection led to the genesis of helpless (H-FST) and non-helpless (NH-FST) mice with a stable phenotype through generations. We characterized these mice for depressive-like behaviors and antidepressant response and compared them to H-TST mice. We also compared their gene expression levels in the prefrontal cortex (PFC) and identified distinct phenotype-associated biological pathways for each model. We next used immunodetection and relative quantification to characterize a specific synaptic impairment in PFC of H-TST and H-FST, suggesting that H-TST and H-FST model different subgroups of MDD with a distinct antidepressant response. Finally, we used two human cohorts to identify a gene-expression based signature that predicts response to duloxetine in women with MDD, using gene expression associated with specific depressive-like behaviors in H-TST and H-FST mice.

## RESULTS

### H-FST and H-TST mice show phenotypic differences in depressive-like behaviors

To generate founders for the selective breeding with a complex genetic background, we performed an eight-way cross of inbred mice (A/J, ABP/Le, BALB/cJ, C57BL/6J, C3H/HeJ, CBA/H, DBA/2J, and SWXL-4). We then used the FST to select animals with the longest and the shortest immobility time and bred them across generations. After ten generations, we observed a 2-fold difference in immobility time between helpless (H-FST) and non-helpless (NH-FST) animals, in both males and females (**Figure 1A**). The percentages of animals reaching scores corresponding to the criteria for selection rose gradually to reach 100% after ten generations for H-FST mice and after fifteen generation for NH-FST mice (**Figure 1A**). Note, we found no difference in motor activity during 45 min in an open field between H-FST and NH-FST mice (not shown). This result suggests that the difference in immobility measured in the FST was not a direct consequence of motor activity modifications.

**Figure 1.**
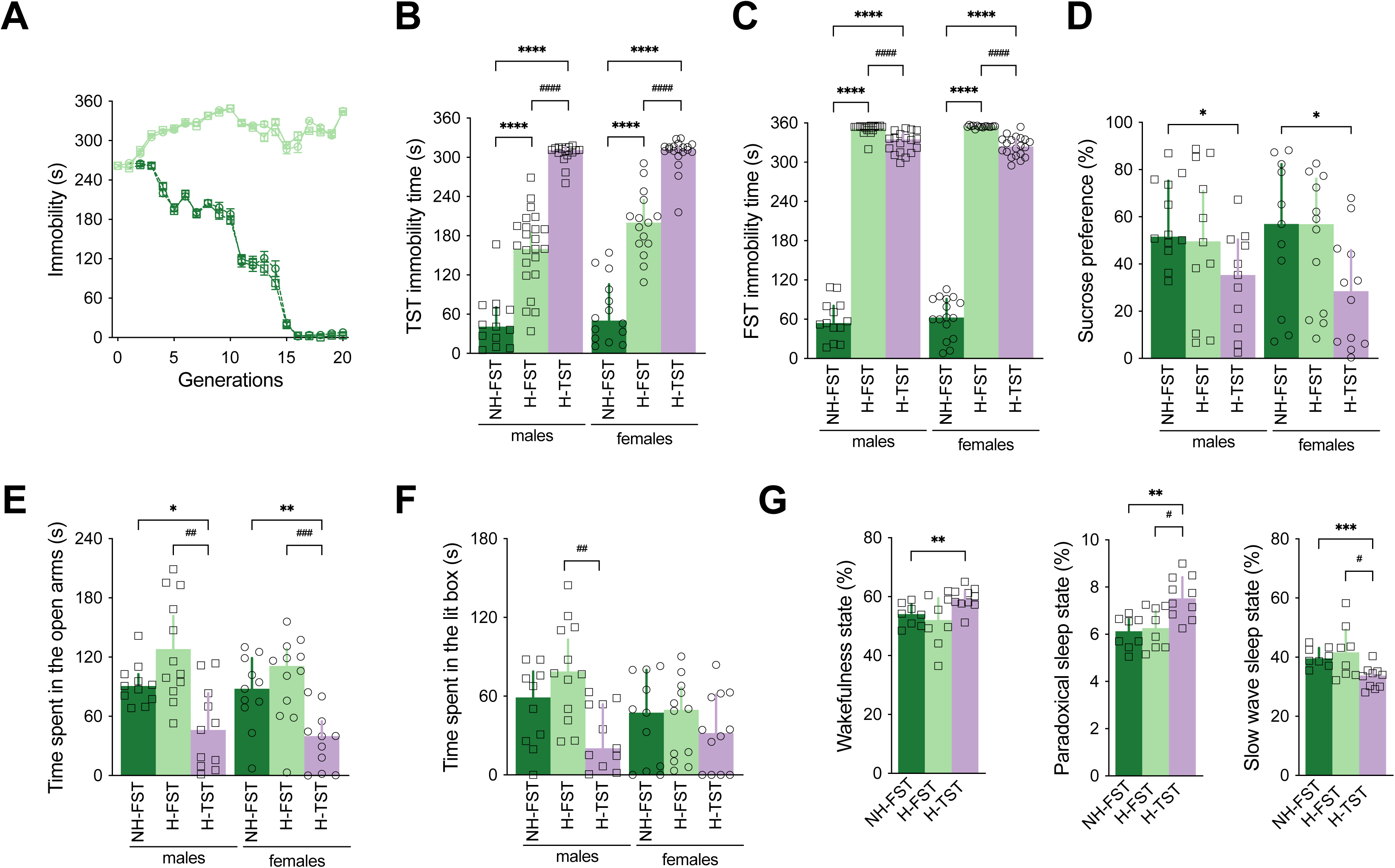
H-FST mice display depressive-like behaviors that differ from those of H-TST mice. **A.** H-FST males and females (light green squares and circles, respectively) show a progressive increase in immobility time in the FST over generations in contrast to NH-FST males and females (dark green squares and circles, respectively) which show a decrease in immobility time. S0 has been estimated using the mean immobility duration of all mice (H-FST and NH-FST). **B, C.** H-FST males (N=23) and females (N=15), and H-TST males (N=18) and females (N=20) display a higher immobility time during the tail suspension test (**B**) and the forced swim test (**C**) than NH-FST males (N=13) and females (N=16), respectively. H-FST mice display a lower immobility in the TST (**B**) and a higher immobility in the FST (**C**) than H-TST mice. **D.** H-FST males (N=11) and females (N=12) do not show a decrease in sucrose preference when compared with NH-FST males (N=12) and females (N=11) respectively, in contrast to H-TST males (N=11) and females (N=12). **E, F.** H-FST mice do not show anxiety-like behavior compared with NH-FST mice. H-TST males (N=11) and females (N=11) spent less time in the open arm of the elevated plus maze, when compared with NH-FST males (N=11) and females (N=11), respectively or to H-FST males (N=11) and females (N=11) (**E**). H-FST males (N=12) and females (N=12) did not spend less time in lit box than NH-FST males (N=10) and females (N=10) (**F**). H-TST males (N=11) but not females (N=12) spent less time in the lit compartment of the dark-light box than H-FST males (**F**). **G**. H-FST females (N=8) do not show any changes in sleep patterns when compared with NH-FST mice (N=8) in contrast to H-TST females (N=10), which spent more time in wakefulness phases and paradoxical sleep and less time in slow wave sleep than NH-FST mice. Scatter plots depict value of all males (squares) and females (circles) tested, and bars represent the median value with interquartile range. Stars symbolize the p-values derived from comparisons with NH-FST mice. Hashtags symbolize the p-values derived from comparisons with H-FST mice. *p<0.05, **p<0.01, ***p<0.001, ****p<0.0001, ^#^p<0.05, ^##^p<0.01, ^###^p<0.001, ^####^p<0.0001.

We then tested H-FST and NH-FST animals for depressive-like behaviors. Using the TST, we showed that helpless males and females displayed a higher immobility time (Mann Whitney test, U_m_=26, p_m_<10^-4^ and U_f_=6, p_f_<10^-4^ for males and females, respectively) when compared with non-helpless NH-FST animals **(Figure 1B**). The immobility time of H-FST mice in the TST was significantly lower than the one of H-TST mice (Mann Whitney test, U_m_=1, p_m_<10^-4^ and U_f_=7, p_f_<10^-4^ for males and females, respectively). In contrast, H-FST mice displayed a higher immobility than H-TST in the FST (Mann Whitney test, U_m_=30, p_m_<10^-4^ and U_f_=4, p_f_<10^-4^ for males and females, respectively), while H-TST displayed a higher immobility than NH-FST (Mann Whitney test, U_m_=0, p_m_<10^-4^ and U_f_=0, p_f_<10^-4^ for males and females, respectively) **(Figure 1C**).

Anhedonia and anxiety are two symptoms frequently reported in patients with MDD, but not in all cases, suggesting they could be used as endophenotypes to determine the most effective treatment methods.^26,27^ We previously showed that H-TST mice displayed anhedonia-like and anxiety-like behaviors.^9^ We thus assessed anhedonia-like behavior using the two-bottle sucrose preference test in the H-FST and NH-FST mice. No difference in sucrose consumption was observed between H-FST and NH-FST mice (Mann Whitney test, U_m_=58, p_m_=0.65 and U_f_=59, p_f_=0.69 for males and females respectively). In contrast, a significant decrease in sucrose consumption was observed for H-TST mice when compared with NH-FST mice (Mann Whitney test, U_m_=25, p_m_=0.01 and U_f_=31, p_f_=0.03 in males and females, respectively). No significant difference was observed between H-TST and H-FST mice (Mann Whitney test, U_m_=40, p_m_=0.19 and U_f_=39, p_f_=0.06 for males and females, respectively) **(Figure 1D**).

We then assessed the anxiety levels using an elevated plus maze **(Figure 1E**) and a light/dark box **(Figure 1F**). In contrast to what was observed for H-TST males and females, NH-FST and H-FST mice spent the same amount of time in the open arms of the elevated plus-maze (**Figure 1E**), suggesting a lower aversion for anxiogenic arms for these mice when compared with H-TST mice (Mann Whitney test, U_m_=26, p_m_=0.02 and U_f_=18, p_f_=0.004 for males and females, respectively). Except for H-TST males which spent less time in the brightly lit box than H-FST males (Mann Whitney test, U=18, p=0.002), no difference was observed between the three lines for females (**Figure 1F**). These results suggest that H-FST mice showed less anxiety-like behavior than H-TST mice.

We previously reported sleep disturbances, a key symptom of MDD, in H-TST mice when compared with NH-TST.^9^ We thus carried out polysomnographic recordings in H-FST and NH-FST mice. We observed no difference in percentage of sleep stages between H-FST and NH-FST mice (**Figure 1G**). Note, H-TST mice spent more time in paradoxical sleep state (Mann Whitney test, H-TST *vs.* NH-FST U=7.5, p=0.002, H-TST *vs.* H-FST U=12.5, p=0.01) and less time in slow wave sleep state (Mann Whitney test, H-TST *vs.* NH-FST U=4.5, p=0.0006, H-TST *vs.* H-FST U=16, p=0.03) than NH-FST and H-FST mice.

In summary, although H-FST mice displayed depressive-like behaviors using the FST and the TST, they showed phenotypic differences when compared with H-TST mice, suggesting the two lines could model the clinical variability observed in individuals with MDD.

### H-FST mice display a reduced immobility in response to antidepressant treatments

One of the major challenges of distinguishing different forms of MDD is the ability to model the heterogeneity of response to treatment that is observed in patients. We previously demonstrated that H-TST mice showed less helplessness when they were treated with LY341495, a selective orthosteric antagonist for the group II metabotropic glutamate receptors (mGluR2/3), than when they were treated with a selective serotonin reuptake inhibitor like fluoxetine.^9^ Therefore to estimate the predictive validity of the H-FST model, we measured the effect of an acute injection of two different selective serotonin reuptake inhibitors, *i.e.,* fluoxetine and escitalopram. An acute intraperitoneal injection of 20 mg/kg fluoxetine in H-FST mice significantly decreased the immobility time in males (Mann Whitney test, U_m_=15, p_m_<10^-4^) and females (Mann Whiney test, U_f=_10, p_f_<10^-4^), with a treatment effect varying between 40% and 50% (**Figure 2A**). Similar results were observed when H-FST mice received an acute intraperitoneal injection of 5 mg/kg escitalopram (Mann Whitney test, U_m_=30, p_m_<10^-4^ and U_f=_12, p_f_<10^-4^ for males and females, respectively) or when they received a daily dose of 10 mg/kg fluoxetine for three weeks (Mann Whitney test, U_m_=17, p_m_=0.03 and U_f_=17, p_f_=0.04 for males and females, respectively; **Figure 2B**). In contrast to what was observed for H-TST mice, a very low treatment effect was observed when H-FST mice received an acute intraperitoneal injection of 3 mg/kg of LY341495 (Mann Whitney test, U_m_=100 p_m_=0.07 and U_f_=54, p_f_=0.01 for males and females, respectively; **Figure 2A**).

**Figure 2.**
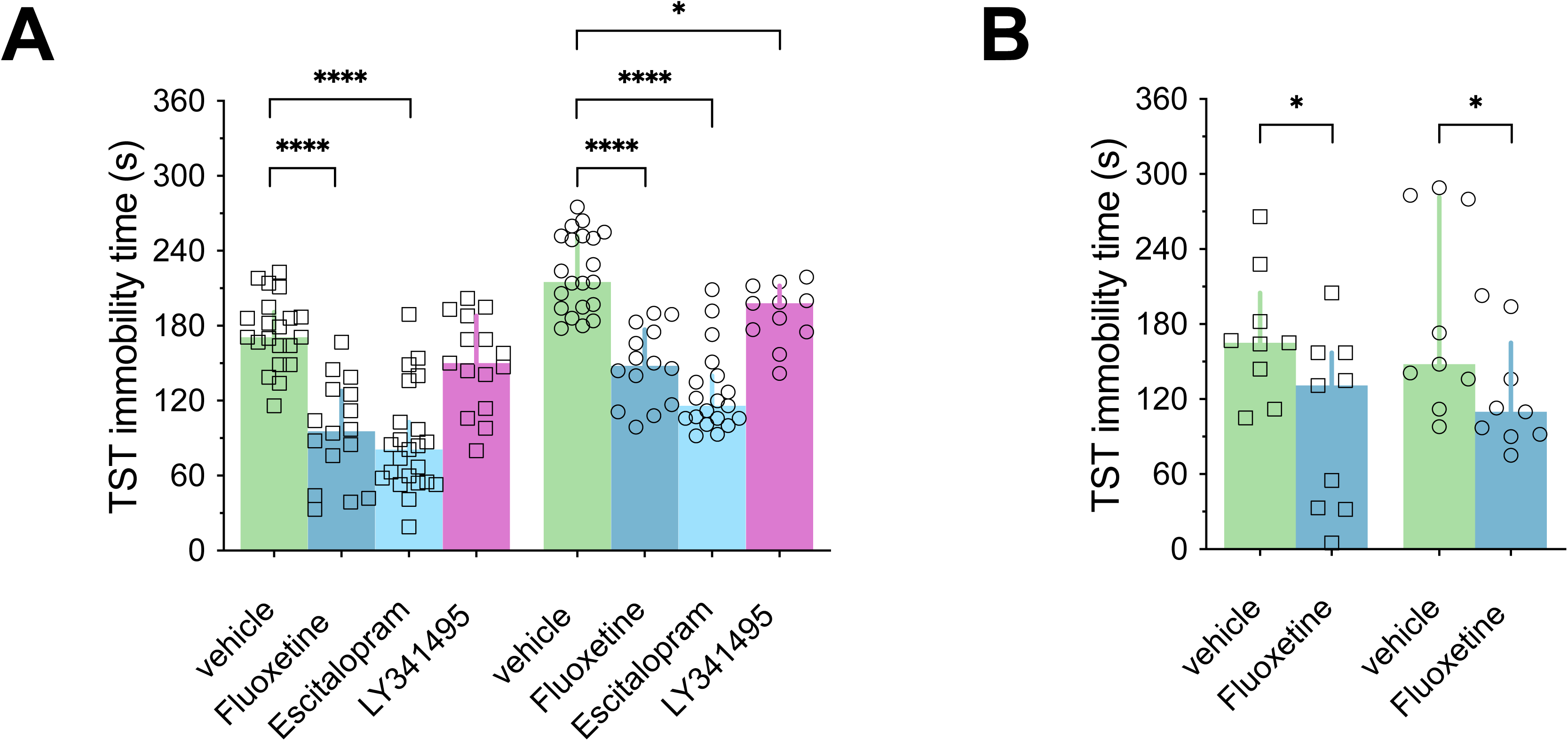
Antidepressant treatments increase the mobility of H-FST males and females in the tail suspension test. **A.** Acute injections: Fluoxetine (20 mg/kg i.p.) substantially increases the mobility of H-FST males (N=15) and females (N=15) in the TST, when compared with a vehicle injection in H-FST males (N=21) and females (N=21). Escitalopram (5 mg/kg i.p) increases the mobility of H-FST males (N=23) and females (N=19) in the TST, when compared with a vehicle injection in H-FST mice. LY341495 (3 mg/kg i.p.) slightly increases the mobility of H-FST females (N=11) but not of males (N=15), when compared with a vehicle injection. **B.** Chronic injections: Fluoxetine (10 mg/kg i.p.) increases the mobility of H-FST males and females (N=9), when compared with vehicle injection (N=9). Scatter plots depict value of all males (squares) and females (circles) tested, and bars represent the median value with interquartile range. *p<0.05, ****p<0.0001.

These observations showed that the selective serotonin reuptake inhibitors were more effective in alleviating the depressive-like behaviors of H-FST mice than glutamate receptor antagonists. This also indicated that H-TST and H-FST mice may model two groups of MDD with different treatment responses.

### H-TST and H-FST mice show specific pattern of gene expression level in prefrontal cortex

In order to characterize the molecular mechanisms underlying the different phenotypes and treatment responses observed in H-TST and H-FST mice, we analyzed the gene expression level in their PFC, a brain region structurally and functionally altered in patients with MDD.^28^ As no phenotypic differences were observed between male and female mice, and given that MDD is more prevalent in females than in males, the gene expression analysis was focused on female mice. Gene filtering resulted in the identification of 14,273 and 14,044 expressed unique genes in PFC of TST and FST mice, respectively. Among them, 13,934 were common between the two mouse lines. Principal components analysis was performed to assess how gene expression profiles could discriminate mouse lines. The three first principal components explained up to fifty percent of observed variance of gene expression in samples **(Figure 3A)**. The first principal component distinguished the selection test, *i.e.*, TST *vs.* FST. H-FST were separated from NH-FST and H-TST from NH-TST according to the second and third principal components, respectively. An unsupervised hierarchical clustering based on their gene expression profile in PFC confirmed the specific gene expression pattern of each mouse line **(Figure 3B)**, TST and FST mice being the most distantly related mouse lines.

**Figure 3.**
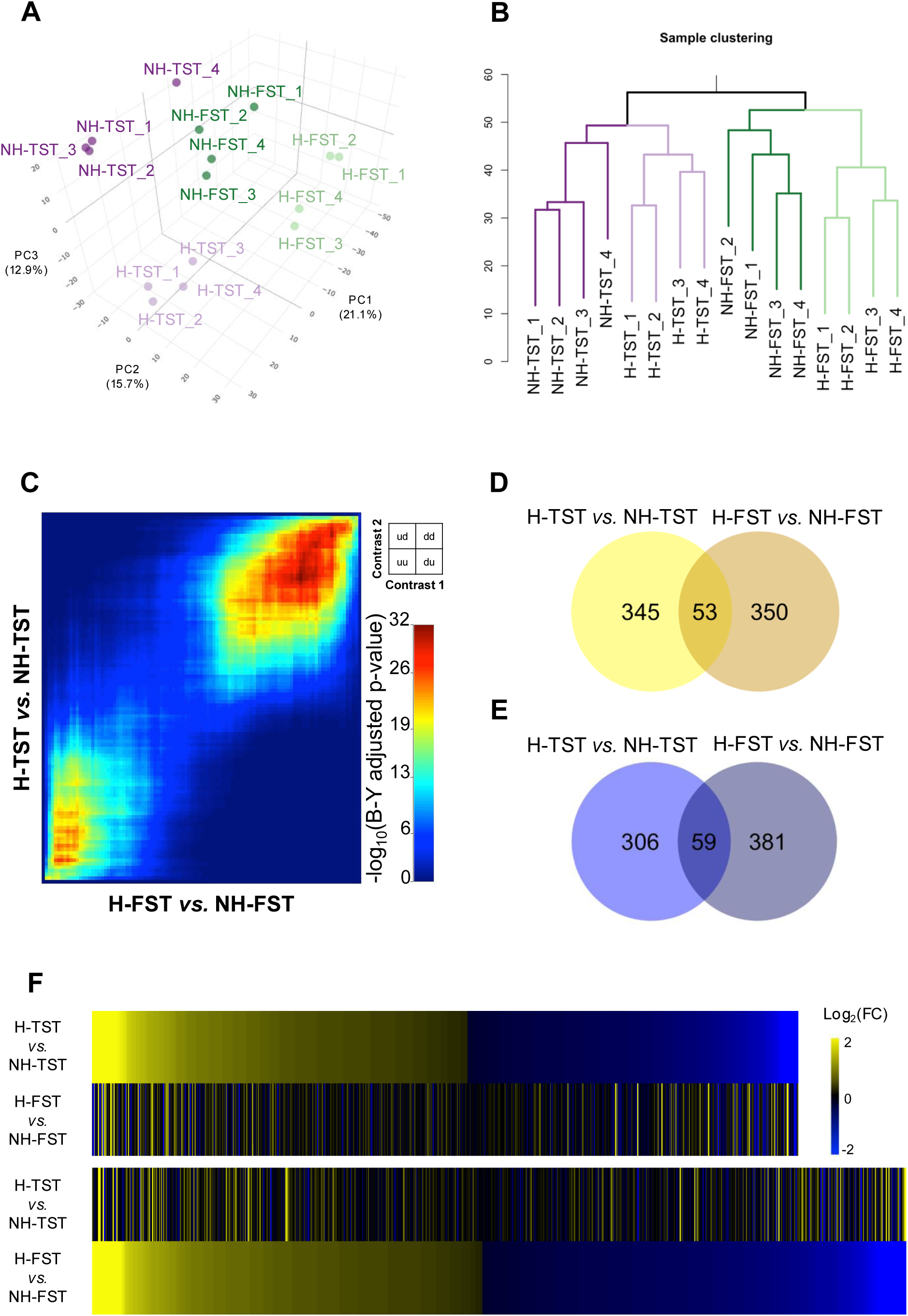
H-TST and H-FST mouse lines show little overlap in their prefrontal cortex gene expression profiles. **A.** 3D plot of sample distribution in the three first principal components (PC) based on their normalized gene expression profile from prefrontal cortex. The variance explained by the component is indicated in brackets. **B.** Sample clustering based on prefrontal cortex gene expression profile using squared Euclidean distance and hierarchical clustering based on average after data normalization. **C.** Heatmap graph of -log_10_ nominal p-values of hypergeometric overlap in transcriptomic changes observed between H-FST *vs.* NH-FST and H-TST *vs.* NH-TST calculated with RRHO technique. d, downregulated genes; u, upregulated genes. **D, E.** Venn diagrams of overexpressed (**D**) and underexpressed (**E**) genes in H-TST and H-FST mice when compared with their non-helpless counterparts. Overexpressed and underexpressed genes are genes with a significant difference (Benjamini-Hochberg adjusted p-value < 0.05) in average expression between helpless and non-helpless mice. **F.** Heatmaps of differentially expressed genes sorted by log_2_(FC) in H-TST *vs.* NH-TST (top) and H-FST *vs.* NH-FST (bottom).

To identify genes that may explain helplessness in H-TST and H-FST mice, differential gene expression between helpless and non-helpless animals per behavioral screening method (H-TST *vs.* NH-TST, H-FST *vs.* NH-FST) was analyzed. We first used unbiased rank-rank hypergeometric overlap (RRHO) analysis to observe that the most differentially expressed genes (DEG) between helpless and non-helpless animals were shared in PFC of H-TST and H-FST mice (RRHO minimal Benjamini-Yekutieli adjusted Fisher exact test, p-value = 10^-32^). **(Figure 3C)**. We also compared H-TST and H-FST PFC differential gene expression pattern with those from other rodent models with depressive-like behaviors induced by genetic or environmental risk factor exposure and for which PFC transcriptome data were available (**Supplementary Figure S1**). RRHO analysis and Pearson’s correlation of gene ranks revealed a good match for up- and downregulation signature between H-TST mice and chronic mild stress mouse model^21^ (RRHO minimal Benjamini-Yekutieli adjusted Fisher exact test, p-value = 2.2 x 10^-9^; Pearson’s rank correlation coefficient = 0.046, p-value = 1.7 x 10^-6^), whereas no positive overlap was observed for H-FST mice (RRHO minimal Benjamini-Yekutieli adjusted p-value = 1.5 x 10^-3^; Pearson’s rank correlation coefficient = -0.046, p-value = 2.7 x 10^-6^). Very few similarities were observed for both models, *i.e.*, H-TST (Pearson’s rank correlation coefficient = -0.019, p-value = 0.051) and H-FST (Pearson’s rank correlation coefficient = -0.013, p-value = 0.19), with animal exposed to chronic social defeat^19^, except for downregulated genes only in the H-TST line (RRHO minimal Benjamini-Yekutieli adjusted p-value = 5.4 x 10^-8^) and the most up- and downregulated genes in H-FST mice (RRHO minimal Benjamini-Yekutieli adjusted p-value = 2.7 x 10^-5^). On the other hand, H-FST mice showed its strongest overlap in up- and down-regulation signature with *Brd1^+/-^*heterozygous mutant female mice (RRHO minimal Benjamini-Yekutieli adjusted p-value = 1.3 x 10^-30^; Pearson’s rank correlation coefficient = 0.12, p-value < 2.2 x 10^-16^), whereas a weaker match with this model was observed only for downregulated genes for H-TST mice (RRHO minimal Benjamini-Yekutieli adjusted p-value = 10^-4^; Pearson’s rank correlation coefficient = 0.013, p-value = 0.18) (**Supplementary Figure S1**).^20^ These comparisons suggest that while few DEG may be shared by some animal models of MDD, most have specificity that could mimic clinical subgroups of MDD.

### Specific biological pathways are affected in PFC of H-TST and H-FST mice

Differential gene expression analysis identified 398 overexpressed genes and 365 underexpressed genes in H-TST mice (**Supplementary Table S1**), and 403 overexpressed genes and 440 underexpressed genes in H-FST mice when compared with the non-helpless mice (**Supplementary Table S2**). Among them, only 112 DEG were common between the two models (53 overexpressed, 59 underexpressed) (**Figures 3D-3E**, **Table 1)**. Interestingly, a single biological pathway was enriched among these genes about the regulation of ERK1 and ERK2 cascade (GO:0070372, adjusted p-value = 0.04). This enriched pathway contained seven genes (*Dusp7, Itgb1bp1, Adam17, Smad4, Ece1, Tlr2, P2ry6*), including four underexpressed genes. Apart from this common mechanism, these results suggest specific gene expression patterns induced by the selection method **(Figure 3F)**.

**Table 1.**
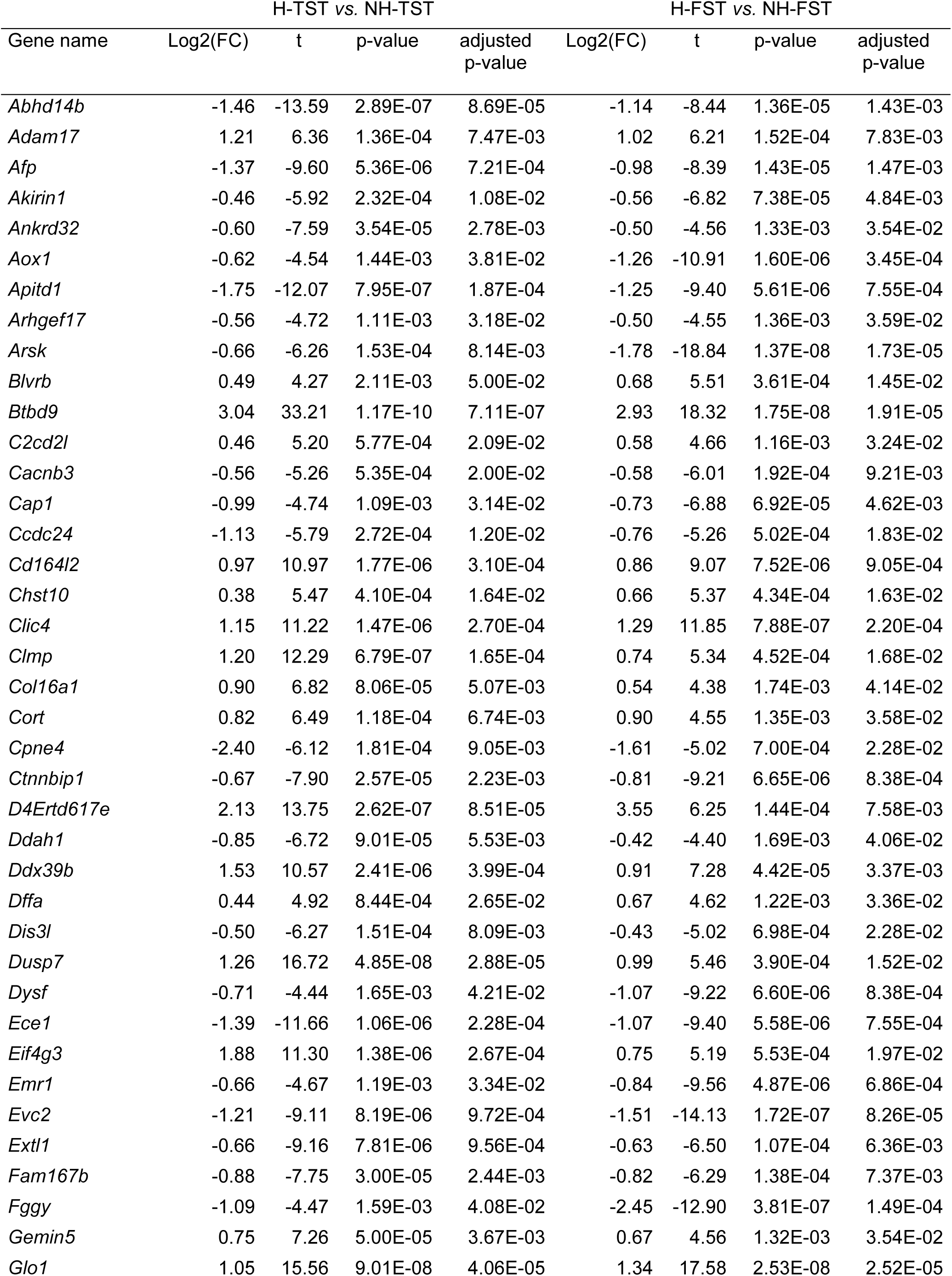
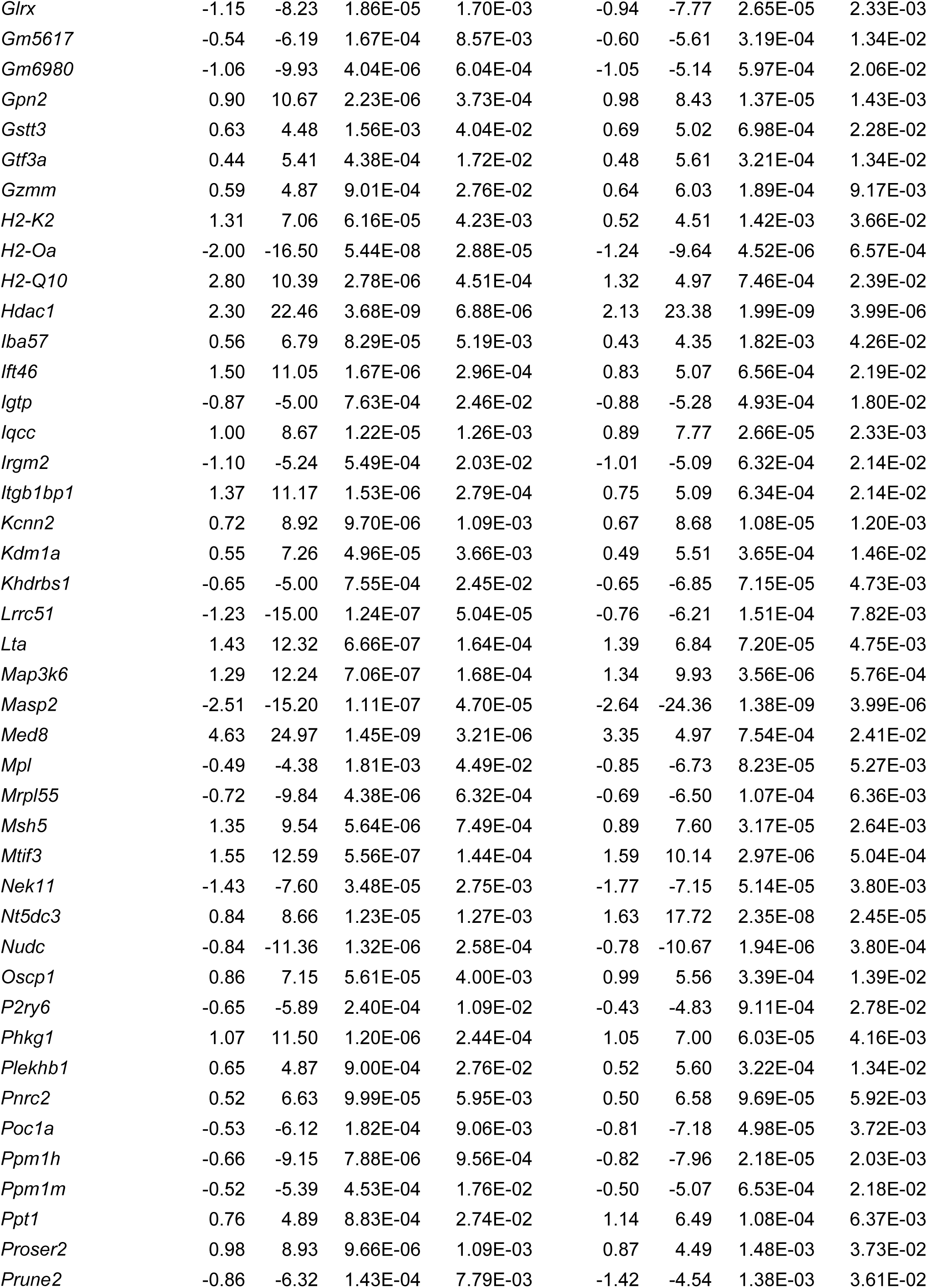
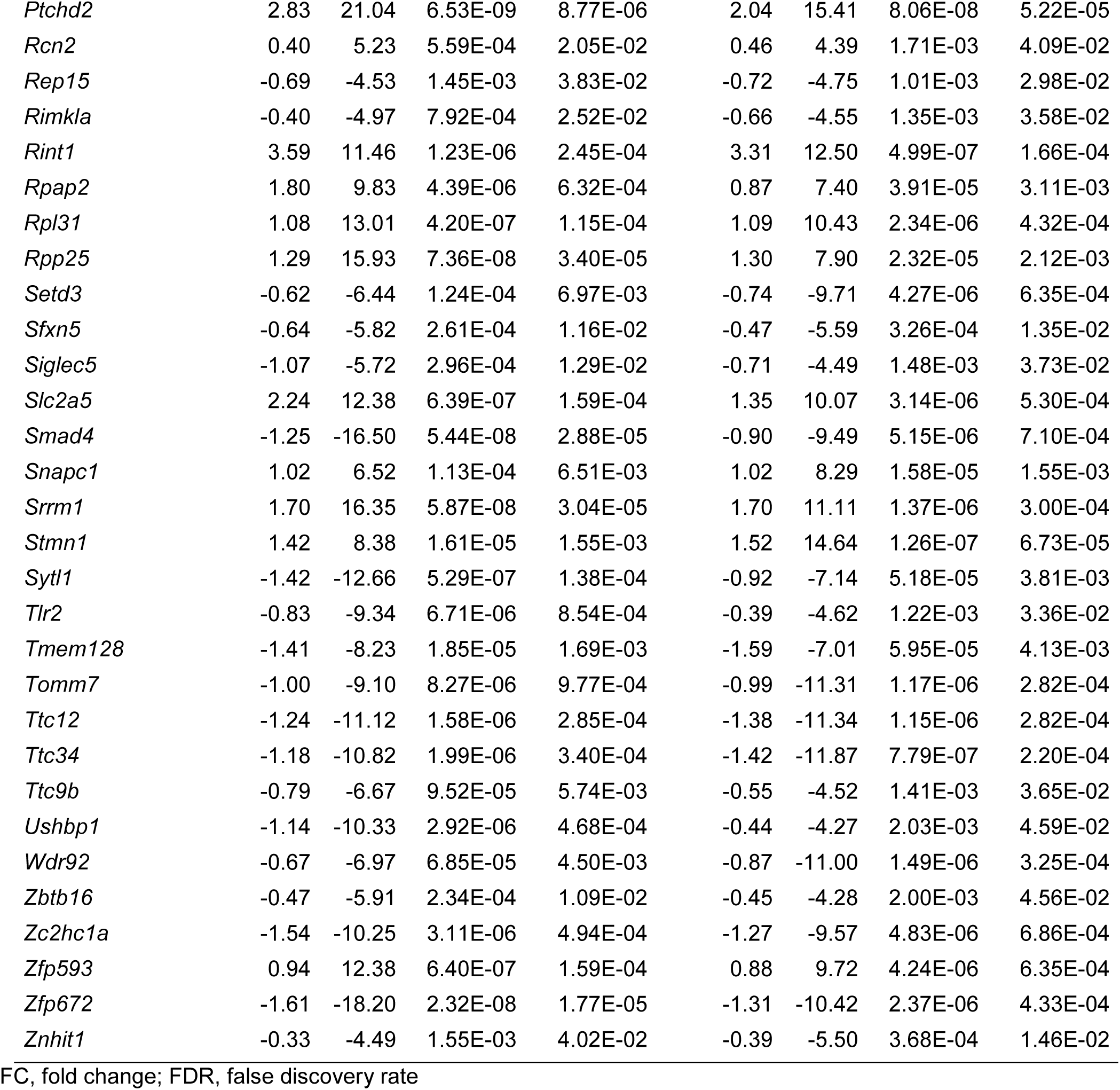
Common differentially expressed genes (Benjamini-Hochberg FDR < 0.05) in H-TST *vs.* NH-TST and H-FST

Biological pathway analysis of DEG confirmed the involvement of mouse line-specific molecular mechanisms **(Figure 4, Supplementary Tables S3 and S4)**. Notably, 100 out of 763 DEG (13.1%) in H-TST were enriched in immunity and inflammation-related processes, constituting the biggest biological pathways cluster of DEG in H-TST mice. On the other hand, 167 out of 843 DEG (19.8%) in H-FST mice were enriched in cell signaling and neurotransmission related processes, constituting the biggest biological pathways cluster of H-FST DEG. Although most of DEG were different between H-TST and H-FST, we noticed that the glutamate receptor signaling pathway and the regulation of response to cytokine stimulus were two biological pathways overrepresented in both mouse lines.

**Figure 4.**
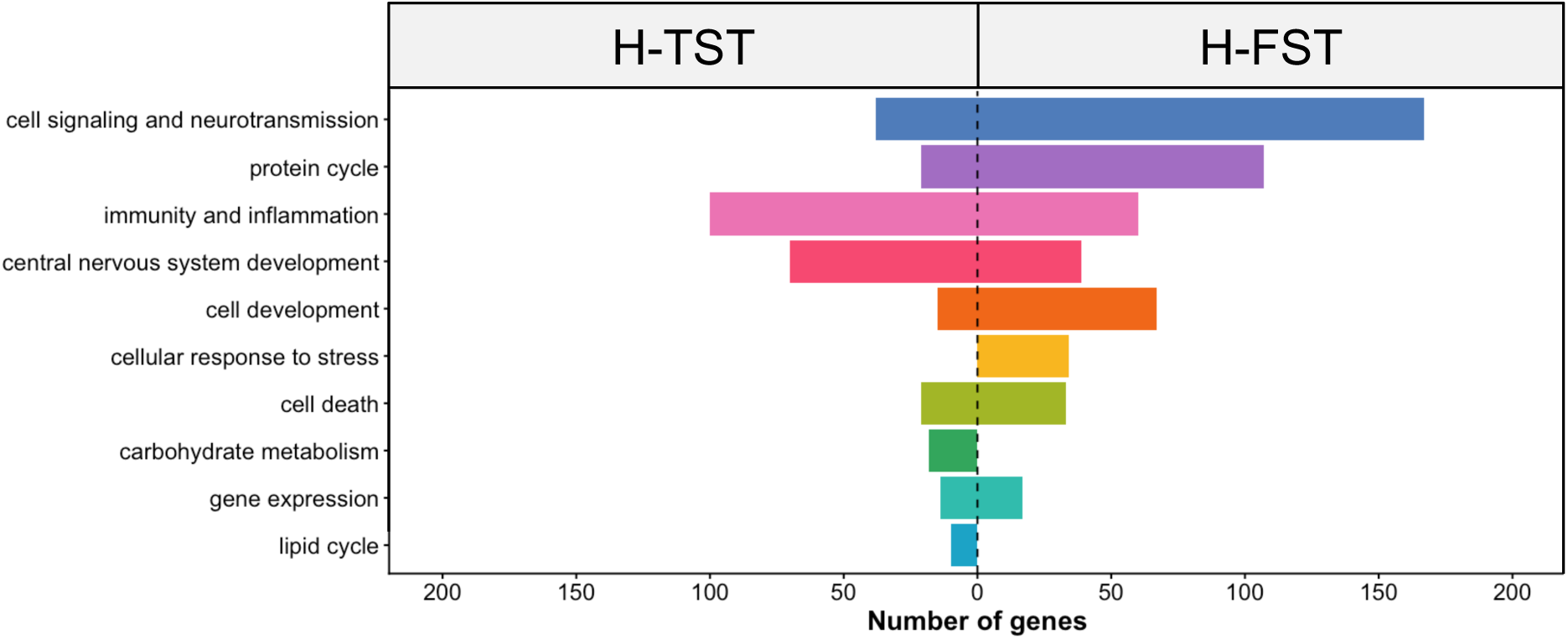
Enriched biological pathways in the prefrontal cortex of H-TST and H-FST mice. Mirror bar charts of clustered enriched biological pathways from differentially expressed genes in H-TST *vs.* NH-TST (left) and H-FST *vs.* NH-FST (right) as listed in the supplementary Table S2. A pathway was qualified as enriched if it contained at least five genes and a Benjamini-Hochberg p-value < 0.05.

### H-TST and H-FST mice showed an imbalance in the equilibrium of the excitatory and inhibitory synapses

Although the glutamate receptor signaling was a biological pathway dysregulated both in H-TST and H-FST, different genes were involved in the two mouse lines (**Supplementary Tables S1 and S2**). To better understand the specificity of each mouse line for this neurotransmitter, we quantified the vesicular glutamate transporter 1 (VGLUT1) and the vesicular GABA transporter (VGAT) in PFC of H-TST and H-FST mice (**Figure 5**). These two transporters are specific biomarkers of excitatory and inhibitory synapses, respectively. Interestingly, we observed a significant higher level of VGLUT1 both in males and females H-TST when compared with NH-TST (**Figure 5B**, Mann Whitney test, U=0, p=0.0002), but no difference was observed for VGAT (**Figure 5C**, Mann Whitney test, U=25, p=0.50). In contrast, a decrease of VGLUT1 and an increase of VGAT was observed for H-FST mice when compared with NH-FST ones (**Figures 5F and 5G**, Mann Whitney test, U_VGLUT1_=2, p_VGLUT1_=0.0006 and U_VGAT_=8, p_VGAT_=0.01). Overall, these results demonstrated that both mouse lines had an imbalance of excitatory and inhibitory synapses in PFC, but in opposite directions (**Figures 5D and 5H**, Mann Whitney test, U_H-TST_=9, p_H-TST_=0.01 and U_H-FST_=6, p_H-FST_=0.005).

**Figure 5.**
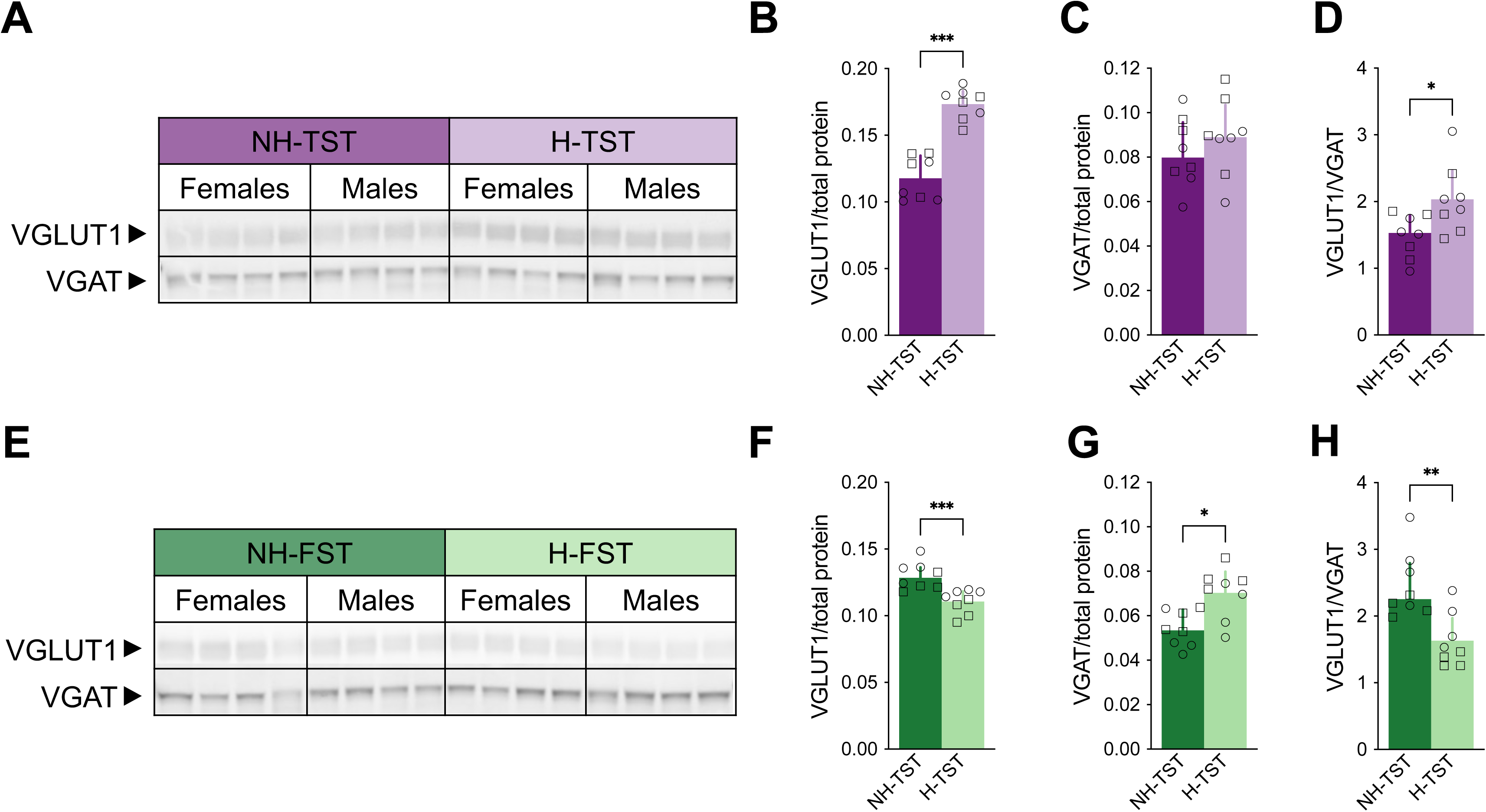
H-TST and H-FST mice showed opposite imbalance of excitatory/inhibitory synaptic markers in prefrontal cortex. **A.** Western blot image (greyscale map) of VGLUT1 and VGAT immunodetected proteins in NH-TST and H-TST mice. **B, C**. Quantification of VGLUT1 (**B**) and VGAT (**C**) proteins relative to total protein amount in NH-TST and H-TST mice. **D.** VGLUT1/VGAT ratio in NH-TST and H-TST mice. **E.** Western blot image (greyscale map) of VGLUT1 and VGAT immunodetected proteins in NH-FST and H-FST mice. **F, G**. Quantification of VGLUT1 (**F**) and VGAT (**G**) proteins relative to total protein amount in NH-FST and H-FST mice. **H.** Scatter plots depict value of all males (squares) and females (circles), and bars represent the median value with interquartile range. *p<0.05, **p<0.01, ***p<0.001.

### Shared genes between human and mouse models identified biomarkers for clinical subgroups of MDD and treatment response

To identify a biological signature distinguishing subgroup of MDD, we performed a RRHO analysis comparing DEG between PFC from H-TST and H-FST mice and DEG between blood of MDD patients with GAD and those without. The human analysis was performed using the GSE98793 dataset that we restricted to women to fit with the mouse transcriptome analysis. The expression level difference was compared for 12,260 genes in common between the mouse and the human analyses. Significant enrichment of downregulated genes both in H-TST female mice and MDD women with GAD was observed (RRHO minimal Benjamini-Yekutieli adjusted Fisher exact test p-value = 4.5 x 10^-4^) **(Figure 6A)**. Among them, 155 genes had a lower expression level (nominal p-value < 0.05) in anxious individuals both in humans and mice (**Figure 6B and Table 2**).

**Figure 6.**
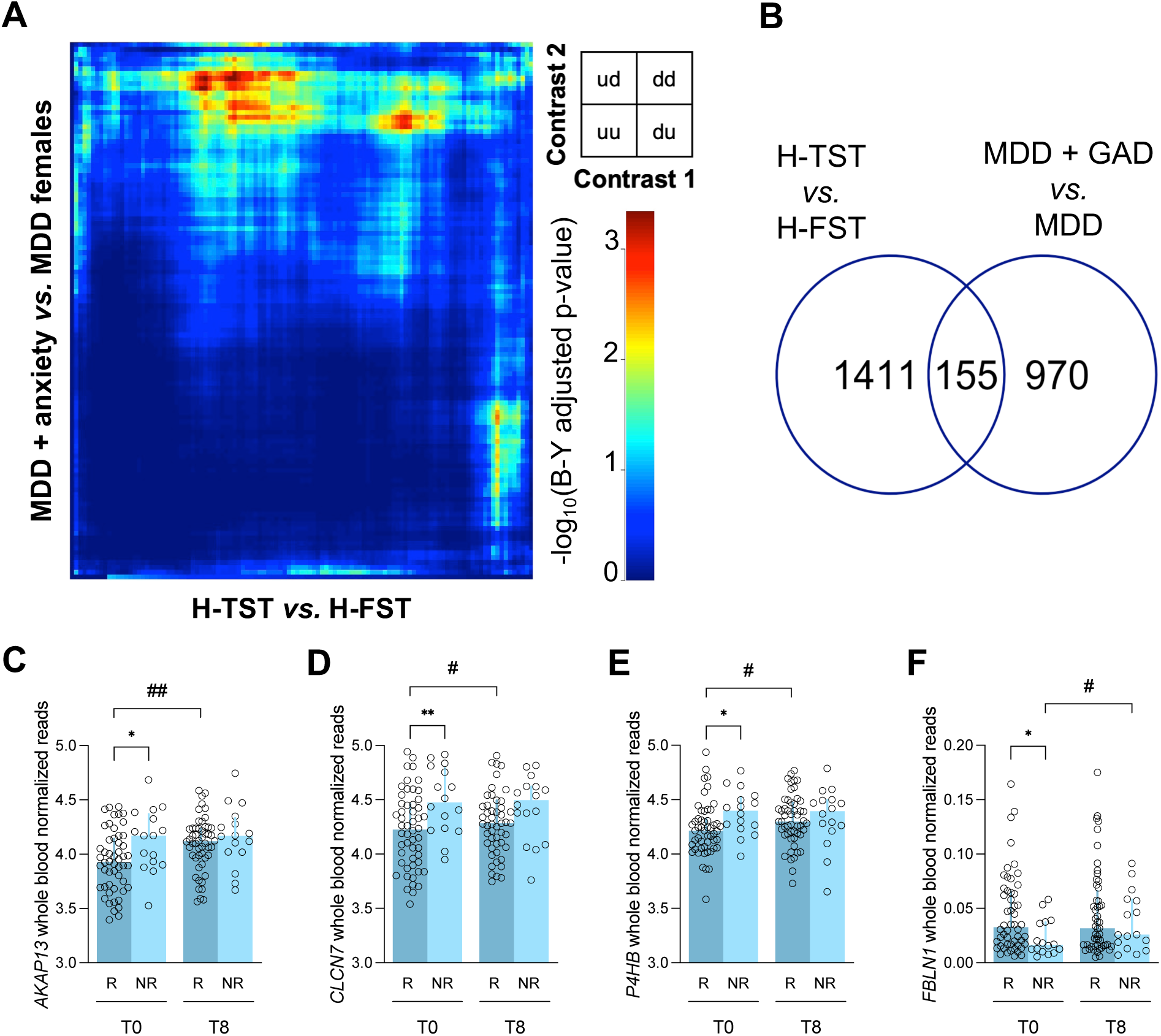
Comparison between genes differentially expressed between H-TST and H-FST mice and those differentially expressed in women with major depressive disorder (MDD). **A.** Heatmap graph of -log_10_ nominal p-values of hypergeometric overlap in transcriptomic changes observed between H-TST *vs.* H-FST and whole blood from women with MDD and generalized anxiety disorder (GAD) *vs.* depressed patients without anxiety calculated with RRHO technique. d, downregulated genes; u, upregulated genes. **B.** Venn Diagram of underexpressed genes (nominal p-value < 0.05) in prefrontal cortex of H-TST mice and whole blood of women with MDD and GAD. **C-F.** Normalized whole blood reads of *AKAP13* (**C**), *CLCN7* (**D**), *P4HB* (**E**) and *FBLN1* (**F**) in women with MDD. Stars symbolize the p-values derived from comparisons between responders (R) and non-responders (NR) to treatment. Hashtags symbolize the p-values derived from comparisons before (T0) and after (T8) eight-week treatment with duloxetine. *p<0.05, **p<0.01, ^#^p<0.05, ^##^p<0.01.

**Table 2.**
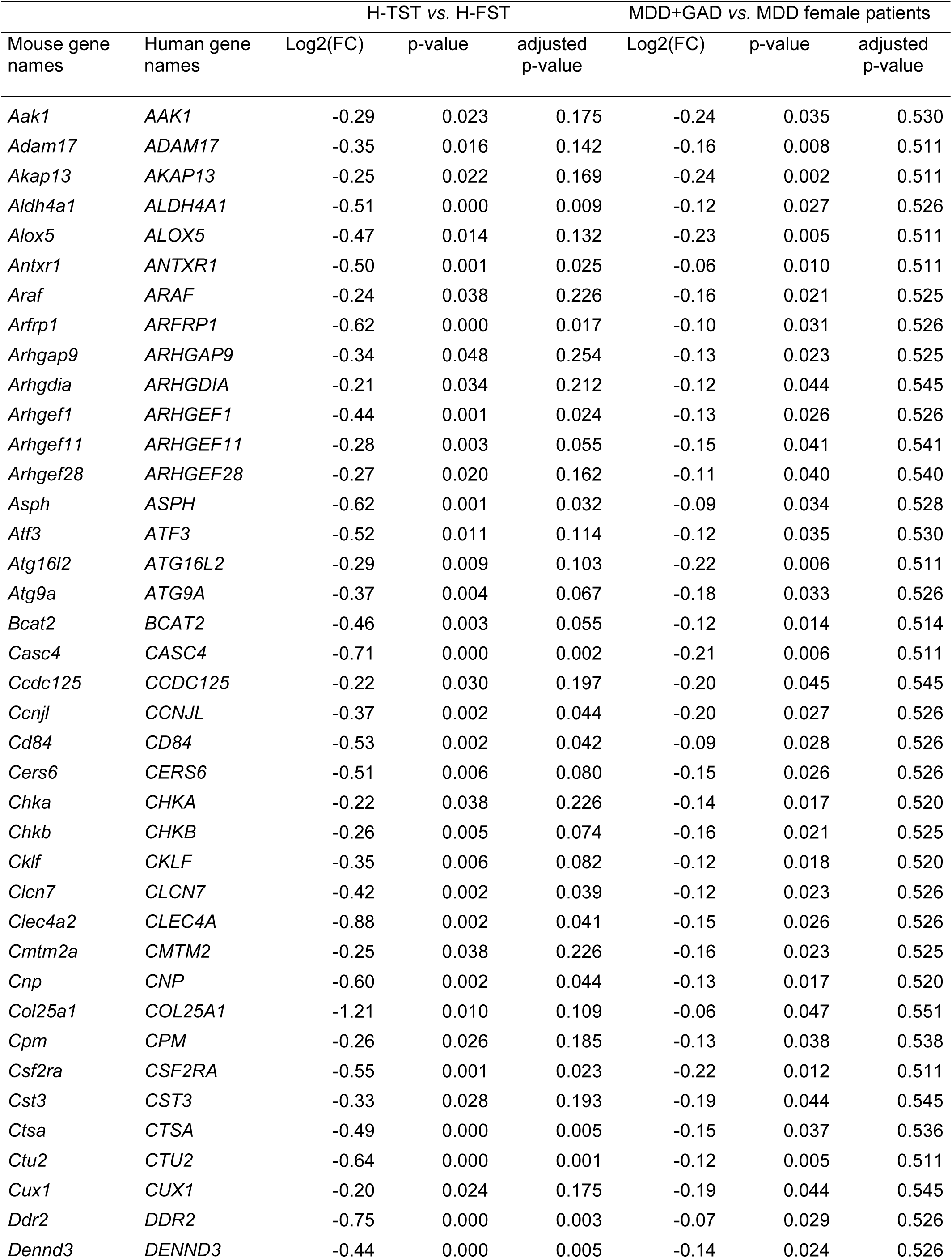
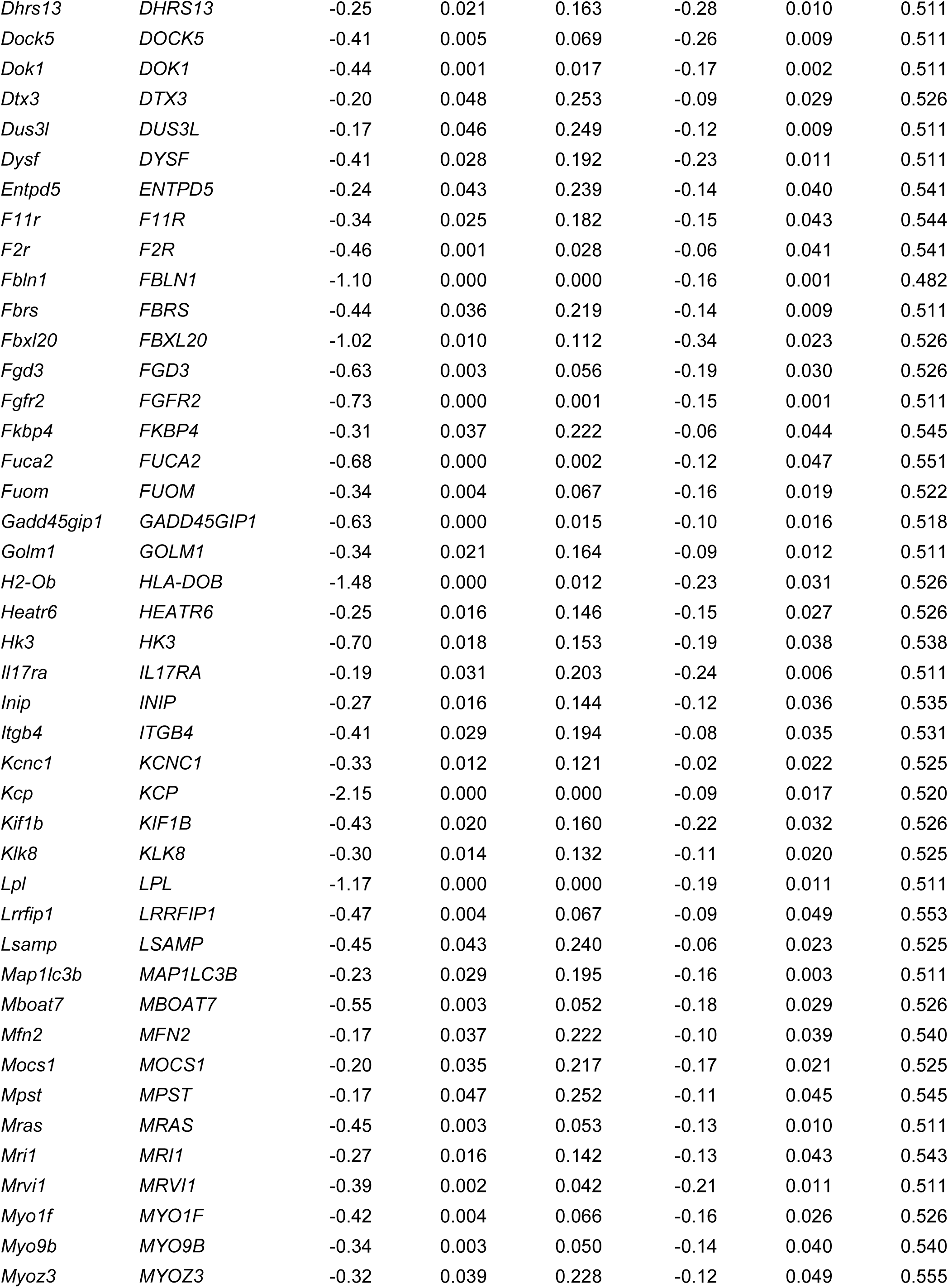
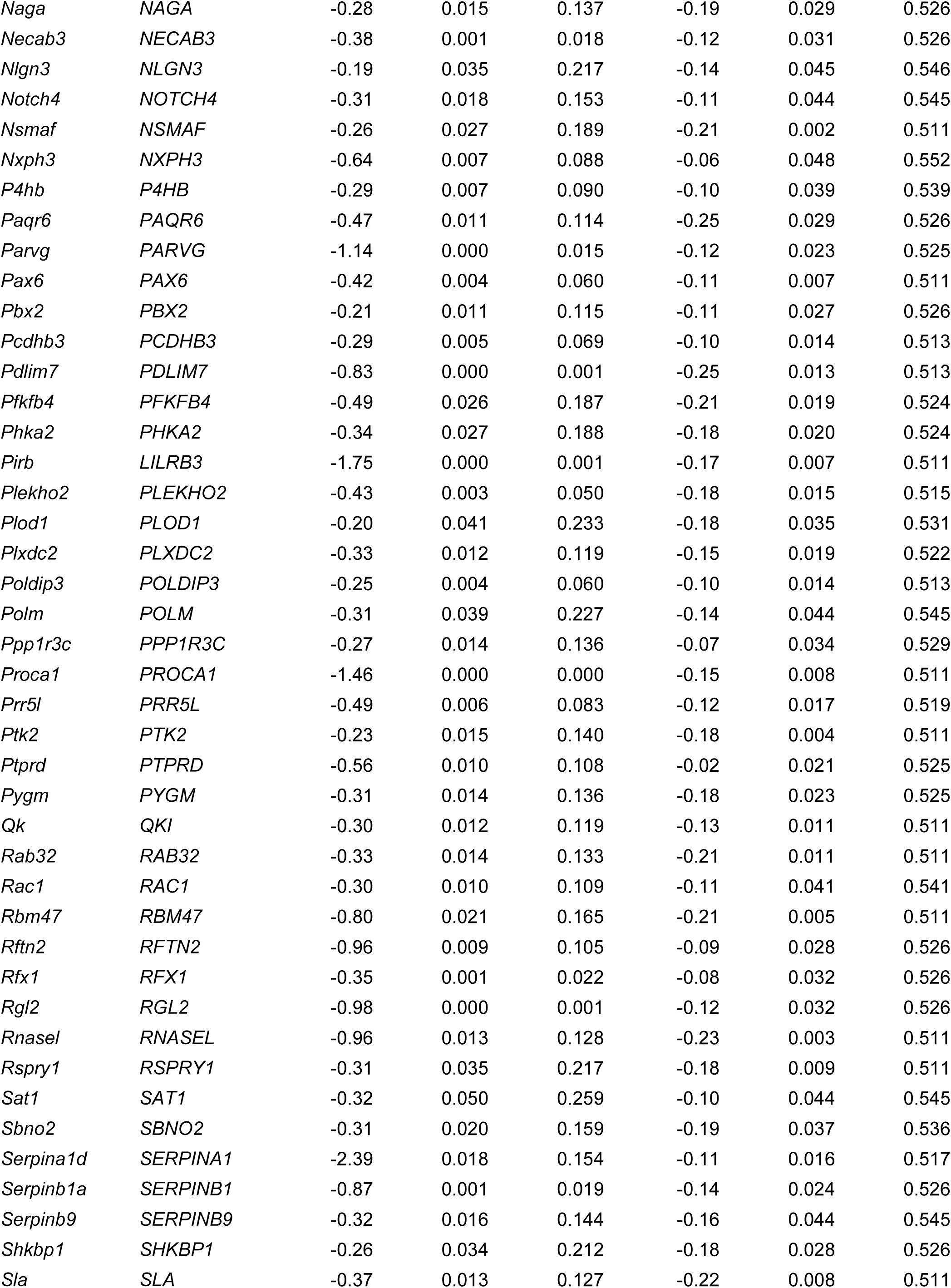
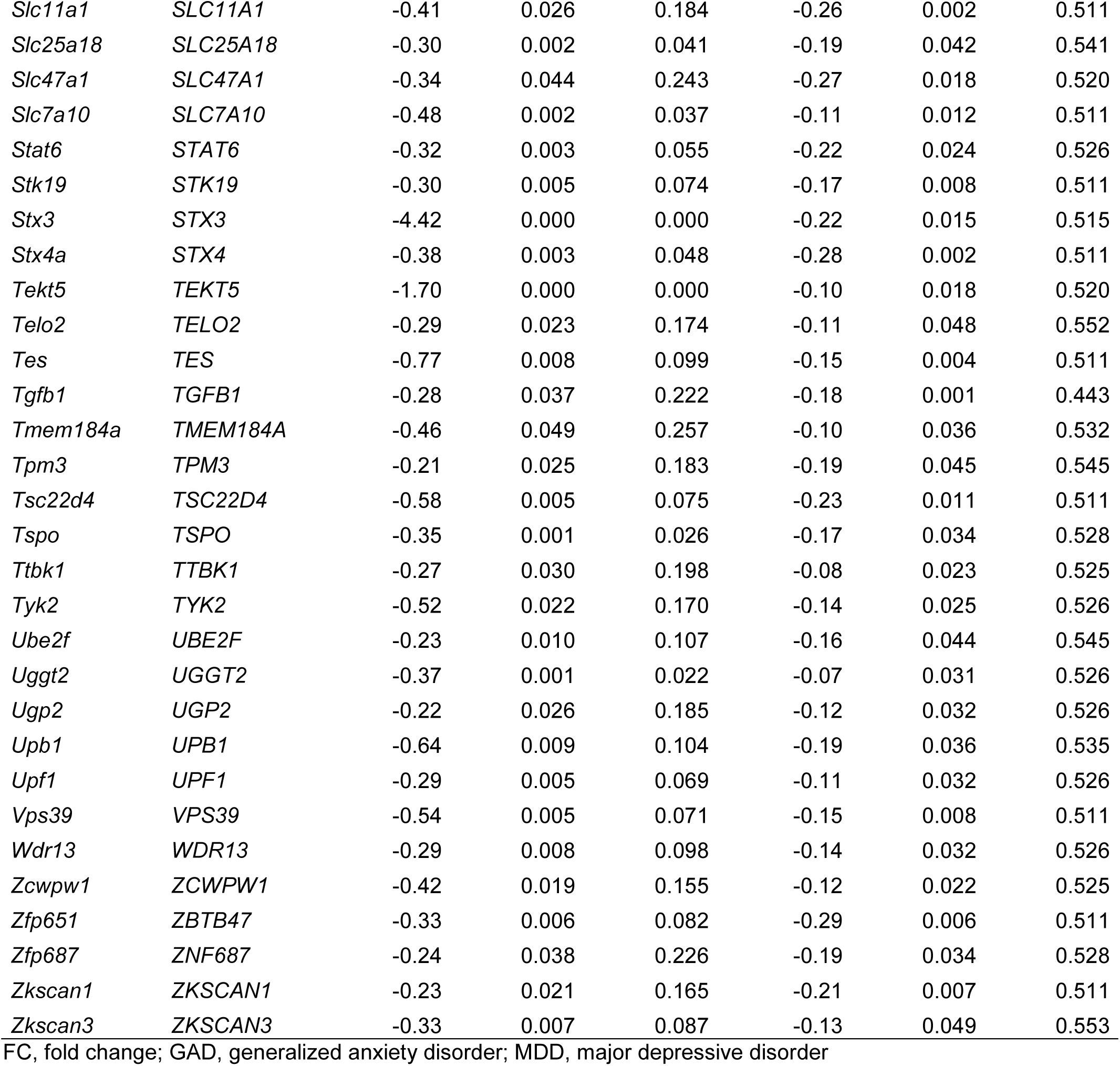
Common underexpressed genes (nominal p-value < 0.05) in prefrontal cortex from H-TST *vs.* H-FST and whole blood from MDD+GAD *vs.* MDD female patients

As serotonin reuptake inhibitors were less efficient on H-TST mice than H-FST ones we assumed that genes under expressed both in H-TST and women with GAD could predict a lower response to standard antidepressant. To test our hypothesis, we used the GSE146446 dataset corresponding to the gene expression level from blood of 237 individuals with a major depressive episode involved in a double-blind clinical trial testing duloxetine at 60 mg daily versus placebo for eight weeks.^25^ Again, we restricted our study to the 68 women who received the duloxetine treatment to remain consistent with our previous analysis. Among the 155 genes that were underexpressed both in H-TST and MDD women with GAD, twelve were differentially expressed (nominal p-value < 0.05) at baseline between women who will respond and those who will not respond to duloxetine after eight weeks of treatment (**Table 3**). Interestingly, three of these genes (*AKAP13*, *CLCN7* and *P4HB*) showed a difference in gene expression level in responders to duloxetine after eight weeks of treatment, and one (*FBLN1*) showed a difference in gene expression level only in MDD women who did not respond to duloxetine treatment (**Figures 6C-6F**). Note, twelve genes in responders and three genes in non-responders showed a difference in gene expression levels after eight weeks of treatment but did not differ at baseline and could be used to predict treatment response only after starting the treatment (**Supplementary Table S5**).

**Table 3.** Genes differentially expressed in prefrontal cortex from H-TST mice, whole blood from women with MDD and generalized anxiety disorders, and whole blood from women with MDD responding to 8-week treatment with duloxetine

| Human gene name | Probe name | T0 |  | T8 |  | Contrast | t | df | p-value |
| --- | --- | --- | --- | --- | --- | --- | --- | --- | --- |
|  |  | R<sup>a</sup> | NR<sup>a</sup> | R<sup>a</sup> | NR<sup>a</sup> |  |  |  |  |
| *AKAP13* | ILMN_2396956 | 3.93 | 4.13 | 4.09 | 4.16 | **NR vs. R (T0)** | **2.45** | **66** | **0.017** |
|  |  |  |  |  |  | T8 vs. T0 (NR) | -0.45 | 15 | 0.657 |
|  |  |  |  |  |  | **T8 vs. T0 (R)** | **-3.44** | **51** | **0.001** |
| *CD84* | ILMN_2049293 | 0.39 | 0.57 | 0.40 | 0.36 | **NR vs. R (T0)** | **2.31** | **66** | **0.024** |
|  |  |  |  |  |  | T8 vs. T0 (NR) | 2.08 | 15 | 0.056 |
|  |  |  |  |  |  | T8 vs. T0 (R) | -0.35 | 51 | 0.725 |
| *CLCN7* | ILMN_1694731 | 4.22 | 4.49 | 4.32 | 4.43 | **NR vs. R (T0)** | **2.74** | **66** | **0.008** |
|  |  |  |  |  |  | T8 vs. T0 (NR) | 0.57 | 15 | 0.576 |
|  |  |  |  |  |  | **T8 vs. T0 (R)** | **-2.10** | **51** | **0.041** |
| *CTU2* | ILMN_2382488 | 0.23 | 0.30 | 0.24 | 0.25 | **NR vs. R (T0)** | **2.21** | **66** | **0.030** |
|  |  |  |  |  |  | T8 vs. T0 (NR) | 0.76 | 15 | 0.462 |
|  |  |  |  |  |  | T8 vs. T0 (R) | -0.34 | 51 | 0.734 |
| *F11R* | ILMN_2270053 | 0.09 | 0.06 | 0.10 | 0.08 | **NR vs. R (T0)** | **-2.40** | **66** | **0.019** |
|  |  |  |  |  |  | T8 vs. T0 (NR) | -1.47 | 15 | 0.162 |
|  |  |  |  |  |  | T8 vs. T0 (R) | -0.27 | 51 | 0.791 |
| *F2R* | ILMN_1742866 | 0.87 | 1.11 | 0.93 | 1.12 | **NR vs. R (T0)** | **2.23** | **66** | **0.029** |
|  |  |  |  |  |  | T8 vs. T0 (NR) | -0.21 | 15 | 0.836 |
|  |  |  |  |  |  | T8 vs. T0 (R) | -1.12 | 51 | 0.268 |
| *FBLN1* | ILMN_1710815 | 0.05 | 0.02 | 0.05 | 0.04 | **NR vs. R (T0)** | **-2.35** | **66** | **0.022** |
|  |  |  |  |  |  | **T8 vs. T0 (NR)** | **-2.16** | **15** | **0.047** |
|  |  |  |  |  |  | T8 vs. T0 (R) | -0.10 | 51 | 0.920 |
| *NLGN3* | ILMN_1759700 | 0.14 | 0.19 | 0.14 | 0.17 | **NR vs. R (T0)** | **2.04** | **66** | **0.045** |
|  |  |  |  |  |  | T8 vs. T0 (NR) | 0.54 | 15 | 0.599 |
|  |  |  |  |  |  | T8 vs. T0 (R) | 0.03 | 51 | 0.975 |
| *P4HB* | ILMN_1719303 | 4.23 | 4.38 | 4.31 | 4.34 | **NR vs. R (T0)** | **2.28** | **66** | **0.026** |
|  |  |  |  |  |  | T8 vs. T0 (NR) | 0.69 | 15 | 0.503 |
|  |  |  |  |  |  | **T8 vs. T0 (R)** | **-2.37** | **51** | **0.022** |
| *PARVG* | ILMN_1695851 | 4.45 | 4.60 | 4.48 | 4.56 | **NR vs. R (T0)** | **2.25** | **66** | **0.028** |
|  |  |  |  |  |  | T8 vs. T0 (NR) | 0.80 | 15 | 0.435 |
|  |  |  |  |  |  | T8 vs. T0 (R) | -0.86 | 51 | 0.392 |
| *TGFB1* | ILMN_2129668 | 0.14 | 0.17 | 0.16 | 0.15 | **NR vs. R (T0)** | **2.13** | **66** | **0.037** |
|  |  |  |  |  |  | T8 vs. T0 (NR) | 0.74 | 15 | 0.470 |
|  |  |  |  |  |  | T8 vs. T0 (R) | -1.08 | 51 | 0.285 |
| *TPM3* | ILMN_1747696 | 0.20 | 0.26 | 0.21 | 0.20 | **NR vs. R (T0)** | **2.03** | **66** | **0.046** |
|  |  |  |  |  |  | T8 vs. T0 (NR) | 1.21 | 15 | 0.244 |
|  |  |  |  |  |  | T8 vs. T0 (R) | -0.69 | 51 | 0.491 |

<sup>a</sup>Mean normalized expression
df, degree of freedom; NR, Non responders; R, Responders; T0, before duloxetine treatment; T8, after eight-week treatment with duloxetine
Significant nominal p-values are indicated in bold

## DISCUSSION

The heterogeneity in antidepressant response among patients with MDD could be explained by the vast heterogeneity in their symptoms, the multifactorial etiology of the disease and its polygenic nature. However, no biological marker is currently available to unravel this complexity and predict response to antidepressants. Here, we generated and characterized a new mouse model of MDD (H-FST) that we compared to another recently published polygenic mouse model of MDD (H-TST).^9^ As observed for H-TST mice, H-FST mice showed helplessness in both the TST and FST, which was stable through generations. Interestingly, H-FST mice did not display anhedonia-like, anxiety-like behaviors and sleep impairments in contrast to H-TST mice. These phenotypic differences between the two mouse lines suggest they may model different subtypes of MDD. Subtyping MDD patients solely based on their clinical profiles to predict antidepressants response appears challenging. However, some symptoms such as anhedonia and anxiety were correlated with depression severity, worse functional outcomes and could predict a poor antidepressants response.^29–34^ Moreover, MDD patients with anhedonia have elevated anxiety, and those with anxious depression are more likely to display anhedonia.^29,35^ Finally, patients with anxious depression exhibit more severe and persistent sleep disturbances than those with non-anxious forms of MDD, where longitudinal studies suggest that sleep disturbances may act as a mediating factor linking anxiety symptoms to the chronic course of depression.^36^ Altogether, anhedonia, anxiety and sleep disturbances, recapitulated in H-TST mice, all have been associated with an increased risk for treatment-resistant depression in patients.^37,38^ Using our behavioral data, we propose the H-TST mouse line as a relevant model of anxious and anhedonic MDD that is less sensitive to classical antidepressants, and the H-FST mouse line as a relevant model of non-anxious and non-anhedonic MDD that is sensitive to selective serotonin reuptake inhibitors (SSRI) antidepressants.

The functional analysis by gene expression comparison between H-TST and H-FST mice demonstrated that our two models shared common profiles but also had specific patterns of differentially expressed genes and pathways in their PFC associated with their specific phenotypes. A great effort is put into data-driven clustering of MDD patients integrating symptoms, treatment response, but also brain functioning and connectivity. As observed for our models, this approach successfully identified subgroups of MDD patients that presented both common and distinct clinical manifestations underpinned by different dysfunctions in brain circuits activity and connectivity. After clustering hundreds of patients based on their resting-state connectivity patterns, Drysdale and colleagues found that hopelessness and anergia was commonly correlated with altered resting-state connectivity between insula, orbitofrontal cortex, ventromedial PFC and multiple subcortical areas in all patients.^39^ However, only the subgroup of patients with anxiety and insomnia symptoms, like H-TST mice, showed a decreased resting-state connectivity between frontal and amygdala regions.^39^ Similarly, Tozzi and colleagues grouped hundreds of patients with depressive and/or anxiety disorders based on their resting-state and task-evoked activity and connectivity within six defined brain circuits.^40^ Again, this work identified alterations in specific circuits associated with some MDD patients presenting anhedonia and anxiety symptoms, with these subgroups showing variability in their response to treatment.^40^ A first group, along with increased anhedonia and anxiety symptoms, had increased task-elicited activity in dorsolateral PFC and dorsal anterior cingulate cortex and was less performant in cognitive control and executive function tasks.^40^ This group had a better improvement than other subtypes after a venlafaxine treatment, a serotonin and norepinephrine recapture inhibitor.^40^ Another group, showed increased anhedonia and anxiety symptoms, accompanied by hyperactivity in stimuli-elicited congruent affects brain circuits.^40^ One hyperactive brain circuit was mobilized by patients’ negative affect evoked by a sad stimulus and comprised pregenual anterior cingulate cortex, anterior insula and amygdala.^40^ The other hyperactive one was mobilized by patients’ positive affect evoked by a happy stimulus and comprised ventromedial PFC and striatum.^40^ Despite the weak transposition of task-evoked functional magnetic resonance imaging in mice, we could use behavioral tasks to assess affective discrimination capability of rodents, and they should be of particular interest to test new therapeutic interventions for MDD.^41^ Altogether, these results suggest that we successfully developed two polygenic models of MDD subgroups that could rely upon different genetic biomarkers and brain functioning. Hence, exploring molecular mechanisms underlying these phenotypes in our models could improve our understanding of this vast heterogeneity observed in both clinical manifestations and antidepressants sensitivity in patients.

Comparison of genes differentially expressed in the H-TST and H-FST mice showed that only 7.5% of them were differentially expressed in both models. Interestingly, “regulation of ERK1 and ERK2 cascade” was the only biological pathway enriched by these genes. In the PFC, the downregulation of the ERK1 and ERK2 signaling cascade is currently one of the most frequent observation associated with DEG involved in pathogenesis, symptomatology and treatment of MDD.^42,43^ Despite the shared complex genetic background in the beginning of the selection process of TST and FST mouse lines to these stressors, the use of these two tests to assess helplessness or non-helplessness led to the generation of four distinct mouse lines with little overlap in the PFC gene expression.

Interestingly, the largest biological pathways differentially expressed in H-TST affect immunity and inflammation. Low-grade inflammation, characterized by increased C reactive protein (CRP), was previously observed in more than a quarter of MDD patients and associated with an increased risk to develop the disease.^44^ Along with this increased CRP, elevated pro-inflammatory cytokines have also been reported in MDD patients.^44^ The inflamed brain hypothesis posits chronic stress disrupts the hypothalamus-pituitary-adrenal axis, leading to an increased permeability of the blood–brain barrier and elevated levels of pro-inflammatory cytokines, which in turn activate microglia and contribute to the development of MDD.^44^ Interestingly, lipopolysaccharide-induced inflammation is associated with worse anhedonia in MDD patients.^45^ Low-grade inflammation also predicts MDD persistence within five years and is associated with a weak response to antidepressants treatments, thus reinforcing the value of H-TST as a model of treatment-resistant depression that could be mediated by neuroinflammation.^46^ Further analysis on neuroinflammation in H-TST and H-FST models is required to conclude.

In contrast to H-TST mice, H-FST mice showed gene expression dysregulation mainly in cell signaling and neurotransmission pathways, and more specifically in the glutamate receptor signaling pathway. Interestingly, we observed an opposite imbalance between excitatory/inhibitory synapses in H-TST and H-FST mice. This observation was consistent with reported variations in glutamate/glutamine levels in frontal regions of patients with mood disorders. On one hand a decreased level of the glutamate/glutamine measure was reported in frontal regions of MDD patients.^47,48^ Note, this deficit was associated with MDD severity and was restored after electroconvulsive therapy.^49,50^ On the other hand, latest meta-analyses from patients with a bipolar disorder found increased glutamate and glutamine levels in frontal regions, independently of their current mood state, manic or depressive.^51,52^ This suggests that our two models may require different treatments to rescue their phenotypes, as recommended for unipolar and bipolar depression. Note, we observed different sensitivity to antidepressants between our models. H-FST mice well responding to SSRI had an excitation/inhibition imbalance towards inhibition in their PFC. On the opposite, H-TST mice well responding to the class II/III glutamatergic antagonist had an excitation/inhibition imbalance towards excitation. Consistently, only MDD patients responding to treatment, whether it was SSRI, ketamine, repetitive transcranial magnetic stimulation or electroconvulsive therapy, were characterized by increased glutamine and glutamate level in prefrontal regions after treatment.^53^ Hence, LY341495, or any other drug targeting class II/III metabotropic glutamate receptors, could be of interest for depressed patients with an excitatory/inhibitory imbalance towards excitation, who are at high risk of not responding to first-line treatments for depression. Moreover, based on their specific molecular impairment, the H-TST and H-FST models should help the identification of biomarkers to stratify MDD patients based on their likelihood to respond to treatment.

Many transcriptome studies conducted in the PFC of MDD patients suggested impairment of synaptic transmission^54^ and more specifically in genes related to glutamate and GABA signaling^55^. At genome level, several single nucleotide polymorphisms (SNP) in glutamatergic genes were associated with increased risk of mood disorders and/or treatment response.^56^ Among them, SNPs within *GRM7,* encoding the glutamatergic metabotropic receptor 7, were associated with early response to citalopram when compared with late response and non-responders in a thousand of MDD patients.^57^ Interestingly, an increased expression of *Grm7* was found in H-TST mice that are weakly sensitive to SSRI, whereas a decreased expression was observed in H-FST mice sensitive to SSRI, suggesting that genes differentially expressed between H-TST and H-FST mice could identify biomarkers of treatment response in humans. We used our PFC transcriptome data to identify translational biomarkers of MDD and antidepressant response. To that end, we found a matching underexpression gene signature between PFC of our anxious *vs.* non anxious helpless models and the whole blood of MDD female patients with *vs.* without comorbid GAD. From this observation, we selected 155 candidate genes to be tested in another cohort of women with MDD responding or not to eight weeks of treatment with duloxetine. This approach led to the identification of four transcripts of interest. *AKAP13* encodes the A-kinase anchor protein 13 involved in downstream signaling of several types of G protein-coupled receptors. Variants of this gene were nominally associated with haloperidol response, characterized by a general decrease in Positive And Negative Symptoms Scale (PANSS) and a specific decrease in the Negative Symptoms PANSS subscale in individuals with a bipolar disorder in a manic state.^58^ This suggests that *AKAP13* could be a relevant translational biomarker of negative symptoms improvement. *CLCN7* encodes the chloride voltage-gated channel 7 and is, just like *AKAP13*, involved in the activation of protein kinase A signaling. Gene and protein product were overexpressed in nucleus accumbens of both MDD and bipolar disorder patients.^59^ Moreover, an expression quantitative trait loci mapped on *CLCN7,* influencing its blood expression level, was associated with postpartum depression, reinforcing its value as a genetic biomarker of depressive episode sensitive to antidepressants.^60^ *P4HB* encodes the beta subunit of prolyl 4-hydroxylase involved in the formation, breakage and rearrangement of protein disulfide bonds. The peripheral blood expression of this gene was increased in postpartum depression and correlated with its severity.^61^ In our analysis, these three genes were all found nominally overexpressed in the PFC of our H-FST model and the whole blood of MDD females without comorbid GAD when compared with their anxious counterparts. Remarkably, these genes levels were also nominally decreased in the whole blood of women responding to duloxetine when compared with resistant at baseline, then increasing with response to duloxetine specifically in responders, pinpointing their relevance for MDD subtyping and duloxetine antidepressant action mechanism. Finally, *FBLN1* encodes the fibulin 1 glycoprotein located in the fibrillar extracellular matrix. Again, this gene was found overexpressed in the PFC of our H-FST model and the whole blood of MDD females without comorbid GAD when compared with their anxious counterparts. This gene was also overexpressed in the whole blood of responders when compared with resistant at baseline and could therefore be a biomarker of good prognosis in MDD patients. Interestingly, four other mRNA studies identified its dysregulation in PFC and blood of stress mice models.^62,63^ A proteomic study in humans also correlated plasma *FBLN1* with the p-factor, a transnosographic psychological score calculated based on externalizing and internalizing symptoms, thus reinforcing its translational value as a biomarker of stress susceptibility and treatment response.^64^

Altogether, the use of bidirectional breeding strategy of mice with a complex genetic background based on their behavioral response to mild stressors successfully reproduced the polygenic nature of MDD, along with its clinical heterogeneity in terms of behavior and antidepressant response. The gene expression study of these models identified a list of candidate biomarkers to predict treatment response in patients with MDD and proposed different pathophysiological mechanisms between responders and non-responders to SSRI treatments. Further use of these models could help the investigation of specific symptoms of the disorder, genetic susceptibility to environmental risk factors and associated brain regions of interest, all towards the identification of disease, treatment response biomarkers and the development of new therapies for a precision psychiatry goal.

## MATERIAL AND METHODS

### Animals

Mice were selectively bred in animal facilities (Faculté de Médecine et de Pharmacie, Lyon 1 University, France) for high or low spontaneous mobility in the FST. The mice used as founders had a broad genetic background resulting from the eight-way cross of A/J, ABP/Le, BALB/cJ, C57BL/6J, C3H/HeJ, CBA/H, DBA/2J, and SWXL-4 mouse strains and had previously been described elsewhere.^9^ These mice were kept on a 7 a.m.-7 p.m. light cycle with food and water *ad libitum*. If not indicated otherwise, behavioral assays were conducted on 10 to 14-week-old mice from generations eight to twenty.

Testing was performed between 9 a.m. and 5 p.m. and was in accordance with the regulations of the local ethical committees CEEA-UCBL 55 (BH 2009-03) and Celyne CEEA-42 (Agreement APAFIS #17130) for the use of experimental animals, as well as the directive 2010/63/EU of the European Parliament and of the Council of 22 September 2010 on the protection of animals used for scientific purposes.

### Selection and breeding

The bidirectional selective breeding was initiated with a starting population of 992 animals (456 males and 436 females) born of 36 breeding pairs. These mice were then selected for helplessness in the TST and the FST. The selection of H-TST and NH-TST mice has previously been described.^9^ Similarly, helplessness was estimated by counting bouts of immobility in a cylinder containing water for FST lines (see below). As for the TST line, we selected the two extreme 10% of all tested mice in a single FST. For the first generation, H-FST were then defined with an immobility time higher than 308 s for males and 310 s for females. Conversely, mice were included in the NH-FST group when their immobility time was lower than 196 s for males and 213 s for females. For the twenty following generations, males and females with the 10% most extreme phenotypes at each generation were then mated together for H-FST on one side, and NH-FST on the other side. Pups were weaned at 21±2 days, and animals were tested in the FST at age 42-50 days.

### Behavioral studies

#### Forced Swim Test

Mice between 7 and 8 weeks of age were plunged individually into a vertical Plexiglas cylinder (25 cm high; 10 cm in diameter) filled with 9 cm deep water (21-23°C) for six minutes. Immobility (*i.e.,* making only minimal movements to keep the head above water or floating) was measured by an observer blind to the condition of the mouse.

#### Tail Suspension Test

The TST was performed with a computerized device. Mice were suspended by the tail with adhesive tape to a hook connected to a strain gauge. This gauge transmitted movements to a computer calculating total duration of immobility during a 6-min test. Mice that climbed up their tail during the test session were excluded from the analysis.

#### Automated locomotor activity test

Exploratory motor activity was monitored in a 50 × 50 cm open-field arena for 45 minutes. Mice, aged 12 weeks, were placed for the test in a dimly lit, silent room. A video camera was placed in the ceiling overlooking the testing area and monitoring four open field cages. The test was performed using a video image processor (Videotrack, View Point, Lyon, France). The total distance travelled by the mice was recorded.

#### Sucrose preference test

Hedonic behavior was assessed using the two-bottle sucrose preference test.^8^ The mice were 14 to 16 weeks old. Consumption of a 2% sucrose solution in one bottle and of water in a second bottle was measured (over a 3-day period) after a 6-day period of habituation with switches between water and sucrose solution as only available fluid. Results are expressed as the percentage of sucrose solution consumed out of the total fluid drunk.

#### Light/dark box

Anxiety was first assessed using the light/dark box test.^8^ This test is based on the innate aversion of rodents to brightly illuminated areas. The light/dark box have two compartments (20 cm large x 20 cm long x 30 cm high) connected by a 6 cm wide by 6 cm height opening located centrally at floor level. The time spent in the brightly 200 lux-lit white (anxiogenic) compartment was recorded during the 5-min testing session using the video-tracking device (Viewpoint).

#### Elevated Plus Maze

Anxiety was also assessed in the elevated plus maze. This maze is a plastic Greek cross placed 50 cm above the floor. The arms were 40 cm long and 6 cm wide. Two opposite arms are surrounded by walls (15 cm high, closed arms), while the two other arms are devoid of such walls (open arms). The mice, aged 13 weeks, were placed for the test in a dimly lit and quiet room. The time spent in the open (anxiogenic) arms over a 5-min period was recorded by the video-tracking device (Viewpoint).

#### Sleep and Wakefulness Analysis

Electrodes for polygraphy sleep monitoring were implanted under general anesthesia as described in detail elsewhere.^9,16^ After a one-week recovery period, female mice (10 to 12-week-old) were placed in a Plexiglas barrel (20 cm in diameter) containing wood chips, food and water *ad libitum*, in a recording chamber for seven days. They were connected to a cable plugged in a rotating connector (Bilaney, Plastics One, Germany) to allow free movements. Unipolar Electroencephalography (EEG) and bipolar electromyography (EMG) signals were amplified (MCP+, Alpha-Omega Engineering, Israel), digitized at 250 Hz, collected with an interface combining the Cambridge Electronic Design (CED) 1401 unit and Spike2 software to analyze off-line using custom scripts (Cambridge Electronic Design Limited, Cambridge, UK). Vigilance states were scored using a 5 s window frame according to standard criteria. After three days of habituation to their cable and new environment, polysomnographic recordings were performed for four days. The analyses were performed using the average of the data obtained after scoring the last two consecutive 24-hour periods.

#### Drug administration

Solutions were prepared daily. Doses always refer to the free bases. Acute treatments were performed on adult mice (10 to 14-week-old). They received an acute intraperitoneal injection with saline, 20 mg/kg fluoxetine HCl (Interchim, Montluçon, France) or 5 mg/kg escitalopram (sigma, France) or 3 mg/kg LY341495 (Bio-Techne, Mineapolis, MN, U.S.A.). All mice were tested in the TST 30 minutes after injection. Chronic treatments were performed in three-month-old mice treated daily, at 10 a.m., with fluoxetine HCl (10 mg/kg i.p.) or saline for three weeks. On day 22, H-FST or H-TST mice were tested (between 1 and 4 p.m.) in the TST.

### Microarray analysis

#### RNA extraction and gene expression quantification

Total RNA was extracted from 20 mg of PFC of four mice per condition. PFC samples were defined as sagittal brain sections of 2 mm of thickness in the frontal cortical region directly posterior to the olfactory bulb. Both left and right sides of the PFC have been collected. FastPrep® homogenizers pulverized PFC through simultaneous beating of specialized Lysing Matrix beads (type D). RNA was isolated on silica-based columns, DNase I digested and eluted with water, using RNeasy Mini Kit following the manufacturer’s instructions (Qiagen, Hilden, Germany). RNA qualities were assessed on an Agilent 2100 Bioanalyzer (Agilent Technologies, Santa Clara, CA, U.S.A.) and RNA had an RNA integrity number higher than 8. Total RNA was quantified using DO 260 nm on a NanoDrop 1000 spectrophotometer (ThermoScientific, Baltimore, MA, USA). Total RNA (50 ng) was amplified and labeled with Cyanine-3 in a round of *in vitro* transcription with Low Input Quick Amp Labeling Kit (Agilent Technologies) following the manufacturer’s protocol. Hybridization was performed following Agilent’s protocol. Briefly, 660 ng of Cy3-labeled cRNA was fragmented and injected onto SurePrint G3 Mouse GE 8×60K Microarrays. Arrays were hybridized 17 h at 65°C with rotation at 10 rpm. After several washes, the arrays were scanned on Agilent C microarray Scanner. The raw data were extracted by Feature Extraction Software v11.5.1 (Agilent Technologies).

#### Differential gene expression analysis

For each sample, gene expression was corrected to reduce background noise and normalized with limma package (v.3.62.2).^17^ Genes were kept in analysis if their Cy3 signal was above background in all samples. Gene filtering led to the identification of 14,273 and 14,044 expressed unique genes respectively in TST and FST mice. Among them, 13,934 were commonly expressed in all mice.

Differential gene expression between helpless and non-helpless animals per screening method (H-TST *vs.* NH-TST, H-FST *vs.* NH-FST) was estimated with functions of the limma package. P-values were adjusted for multiple comparison using a false discovery rate (FDR) estimation based on the Benjamini-Hochberg (B-H) method.^18^ A gene was considered differentially expressed if its B-H adjusted p-value was below 0.05.

Other gene expression datasets for mouse models comparison can be accessed at GSE81672^19^ and GSE150265^20^. Raw data from chronic mild stress mouse model^21^ and transcript per million data from *Brd1^+/-^* heterozygous mutant female mice were kindly provided by authors. Raw reads from chronic mild stress were transformed and quantile normalized and differential gene expression between mouse models *vs.* control mice was estimated with functions of the limma package. Raw counts from chronic social defeat mice were transformed and normalized using trimmed mean of M values and differential gene expression between mouse models *vs.* control mice was estimated with functions of the edgeR package (v. 4.4.2).

#### Rank-rank hypergeometric overlap (RRHO)

Inversed log_10_-transformed nominal p-values were ranked based on their decreasing value multiplied by the sign of the log_2_(fold change) between helpless and non-helpless animals. Sorted gene lists were compared through rank-rank hypergeometric overlap (RRHO) analysis using the RRHO package (v. 1.46.0).

#### Gene ontology analysis

Differentially expressed genes were annotated with their biological pathways, and each proportion of differentially expressed genes in a given biological pathway were tested for over-representation using hypergeometric distribution to calculate the associated p-value with functions from clusterProfiler package (v.4.14.6).^22,23^ A pathway was significantly enriched if it contained at least five genes and a B-H-adjusted p-value bellow 0.05. Gene ontology (GO) terms were then subclustered based on their semantic similarity, reduced in parent terms with functions from rrvgo package (v.1.18.0) and broad clusters were manually assigned.

#### Human datasets

Whole blood gene expression from 96 individuals, 64 of whom were diagnosed with MDD including 32 with comorbid generalized anxiety disorder (GAD) were available on Gene Expression Omnibus (GEO) accession GSE98793.^24^ In addition, whole blood gene expression from 406 individuals with MDD treated with duloxetine, citalopram or a placebo and for whom antidepressant response was assessed after eight weeks of treatment was collected at GEO accession GSE146446.^25^ Before analysis, gene expression data from GSE102556 and GSE146446 was transformed and normalized using the limma package.

### Western Blots

#### Protein extraction

Four prefrontal cortices per condition (mouse lines, helplessness and sex) were mechanically grinded in 50 µL of lysis buffer, containing 50 mM Tris-HCl, 100 mM NaCl, 1% Triton X100, 5 mM Ethylene-Diamine-Tetra-Acetic acid (EDTA), 1 mM Phenylmethylsulfonyl fluoride (PMSF), and 1 mini-tablet of proteases and phosphatases inhibitors cocktail (Ref. A32961, Thermo Fisher Scientific, Waltham, MA, U.S.A.). Samples were then centrifuged for 10 minutes at 10,000 x g at +4 °C. Supernatant S1 proteins were collected and dosed using Bradford assay.

#### Protein immunodetection and relative quantification

For the vesicular glutamate and the gamma-amino butyric acid (GABA) transporters (VGLUT1 and VGAT, respectively) immunodetections, 20 µg of non-heated S1 protein samples were resolved in NuPAGE™ Bis-Tris 10% polyacrylamide gels (Thermo Fisher Scientific) with 1X NuPAGE™ MOPS running buffer (Thermo Fisher Scientific).

Proteins were transferred onto a low fluorescence polyvinylidene fluoride membrane overnight at +4 °C in 1X tris-glycine added with 10% methanol. For protein level normalization, total proteins were detected with No-Stain™ labelling kit (Thermo Fisher Scientific) as described by the manufacturer. Protein immunodetection was performed following the Azure fluorescent Western Blot workflow (Azure Biosystem, Dublin, CA, U.S.A.). As primary antibodies, we used a monoclonal mouse antibody against VGLUT1, clone 68B7 (Ref. 135 011, Synaptic Systems GmbH, Göttingen, Germany) and a polyclonal rabbit antibody against VGAT (Ref. 14471-1-AP, Proteintech, Rosemond, IL, U.S.A.) at a 1:5,000 dilution in 1X Azure fluorescent blot blocking buffer (Ref. AC2190, Azure Biosystem) and incubated for 1.5 hours at room temperature. As secondary antibodies, we used a goat anti-mouse IR800 (Ref. AC2135, Azure Biosystem) and a goat anti-rabbit IR700 (Ref. AC2128, Azure Biosystem) at a 1:10,000 dilution in 1X Azure fluorescent blot blocking buffer for 1 hour at room temperature.

### Statistical analysis

Statistical analysis was conducted with R v.4.4.2. All normally distributed variables were expressed as mean ± standard deviation (SD) and comparisons between conditions were examined by t-tests. Not normally distributed variables were expressed as median and interquartile range (IQR) and between-group comparisons were analyzed by Mann–Whitney test.

## Data availability

All data and scripts are available on Figshare or upon request.

## Acknowledgments

This work was supported by the French government managed by the *Agence National de la Recherche* under the 2012 generic program (ANR-12-SAMA-0012-01, project DEPSOM) and the France 2030 program (ANR-22-EXPR-0004). It was also supported by the *Fondation de France* under references 00081240 and 00091264. This work also received financial support from the *Institut National pour la Santé et la Recherche Médicale* (Inserm), the *Centre National de la Recherche Scientifique* (CNRS), and the *Université Paris Est-Créteil* (UPEC). We acknowledge the SFR Santé Lyon-Est (UAR3453 CNRS, US7 Inserm, UCBL) facilities, particularly the ALECS Conventionnel platform. We thank the animal care staff for their technical support, and Stéphane Marinesco and Fabienne Rajas for enabling the housing of mouse lines in Lyon and access to the ALECS facilities for behavioral and related experimental studies. We also acknowledge Université Claude Bernard Lyon 1 for institutional support and hosting of the animal facilities. We would like to thank Benoît Martin for providing us with the mouse strains used to generate the founders and for his advices on selective breeding.

## Authors contributions

CA performed molecular biology and biochemical experiments and transcriptome analyses. MEY, JMV and SJ designed the study, supervised the project and contributed to the interpretation of the data. MEY and JMV generated mouse lines by selective breeding with the help of MD. MEY conducted mouse behavioral analyses and pharmacological studies, with the help of SA for sleep exploration. VL helped with molecular biology experiments. CA and SJ wrote the article with the help of MEY and JMV.

